# Expression weighted cell type enrichments reveal genetic and cellular nature of major brain disorders

**DOI:** 10.1101/032680

**Authors:** Nathan G. Skene, Seth G.N. Grant

## Abstract

The cell types that trigger the primary pathology in many brain diseases remain largely unknown. One route to understanding the primary pathological cell type for a particular disease is to identify the cells expressing susceptibility genes. Although this is straightforward for monogenic conditions where the causative mutation may alter expression of a cell type specific marker, methods are required for the common polygenic disorders. We developed the Expression Weighted Cell Type Enrichment (EWCE) method that uses single cell transcriptomes to generate the probability distribution associated with a gene list having an average level of expression within a cell type. Following validation, we applied EWCE to human genetic data from cases of epilepsy, Schizophrenia, Autism, Intellectual Disability, Alzheimer’s disease, Multiple Sclerosis and anxiety disorders. Genetic susceptibility primarily affected microglia in Alzheimer’s and Multiple Sclerosis; was shared between interneurons and pyramidal neurons in Autism and Schizophrenia; while intellectual disabilities and epilepsy were attributable to a range of cell-types, with the strongest enrichment in interneurons. We hypothesized that the primary cell type pathology could trigger secondary changes in other cell types and these could be detected by applying EWCE to transcriptome data from diseased tissue. In Autism, Schizophrenia and Alzheimer’s disease we find evidence of pathological changes in all of the major brain cell types. These findings give novel insight into the cellular origins and progression in common brain disorders. The methods can be applied to any tissue and disorder and have applications in validating mouse models.

## Introduction

The brain has a highly complex cellular architecture characterized by a diverse set of cell types that are highly interconnected. Identifying the cell types involved with the pathogenesis of disease is particularly challenging in heterogeneous tissues where cell types are often poorly defined. In the majority of brain disorders evidence exists for changes affecting multiple cell types. It has proven problematic to determine which cells are associated with the primary disease pathology and which are altered as secondary “reactive” responses. Genomic technologies have contributed important mechanistic insights into the primary genetic basis of pathogenesis through studies of mutations and variants that increase disease susceptibility. The recent availability of single cell transcriptomes (SCT) from brain tissue[1; 2; 3; 4; 5; 6] offer the opportunity to ask which cells express these susceptibility genes, and thereby identify the primary cellular pathology. Since changes in primary and secondary cell types will be reflected in transcriptomic studies of diseased tissue, SCTs could also be used to identify these two classes of cell types.

A method known as Population Specific Expression Analysis (PSEA)[7] was previously developed with the goal of extracting cell-type data from disease transcriptomes and has been successfully used to determine the cellular pathology in Huntington’s and Parkinson’s disease[8]. A limitation with this method was the dependence on predefined sets of cellular genes that are specific to particular cell types. As the method depended on linear modeling, it could not be applied to unquantified gene sets thereby limiting its applicability for use in gene expression studies. To overcome these limitations, we have developed the Expression Weighted Cell-type Enrichment (EWCE) method that statistically evaluates whether a set of genes has higher expression within a particular cell type than can be reasonably expected by chance. For the purposes of the current study, the cell type expression profiles are defined using data from single cell RNA-sequencing. The method can be applied to any experimental source that generates a list of genes, including but not limited to genetic, transcriptomic and proteomic studies. The workflow for two such experimental designs are shown in Figure 1a alongside data from an example gene set, which demonstrates the principles of the methods in Figure 1b.

**Figure 1.**
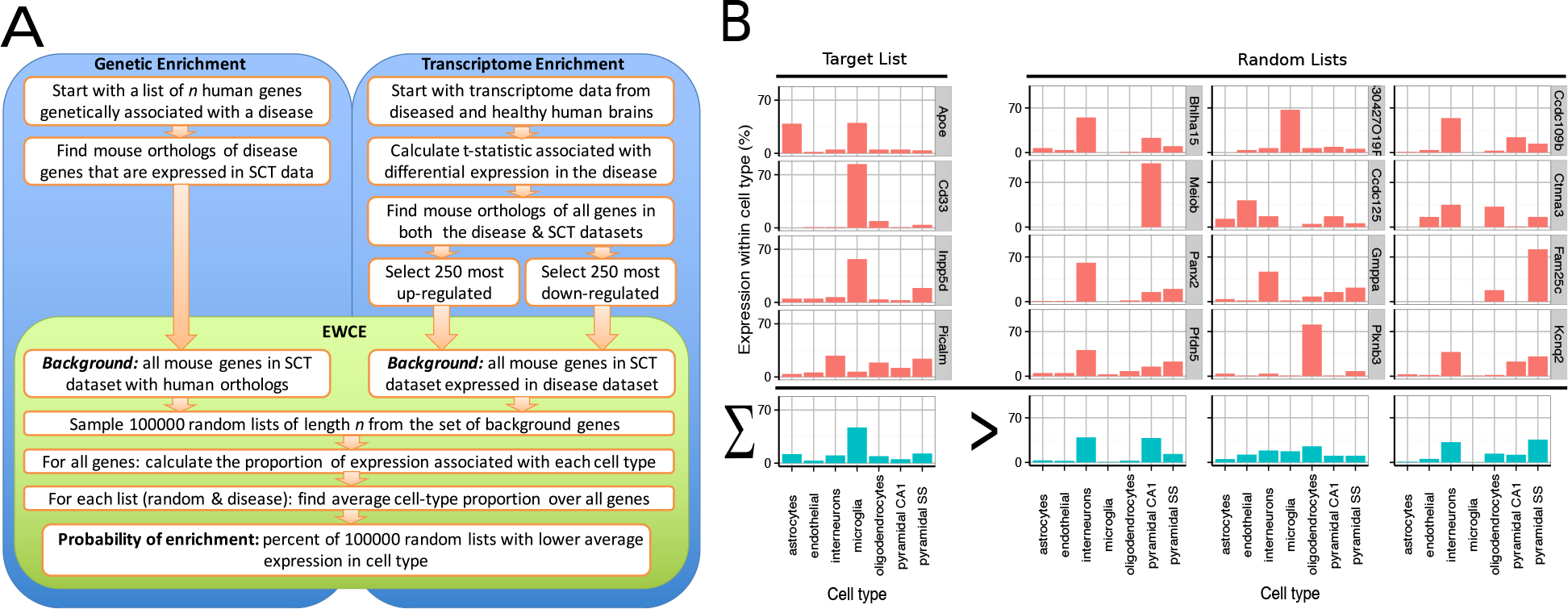
Depiction of the Expression Weighted Cell-type Enrichment method. *(A) Flow diagram showing the steps involved in going from either genetic or transcriptional data to a probability of enrichment.* *(B) Demonstration of the principles ofEWCE method using four Alzheimer’s genes (target list) and three randomly generated lists. In the left most column, the cell type expression proportions for the target list is shown. The row with blue bars shows the average over the genes shown above. All four of the target genes are not specific marker genes for microglia; Apoefor instance also has high expression in astrocytes. Nonetheless, when averaged together, there is a higher mean expression in microglia than in the averaged random list.*

The EWCE method has been enabled by recent studies that have dramatically increased the availability of single cell transcriptomes from brain tissue[1; 2; 3; 4; 5; 6]. As many of these studies selected the cells through unbiased sampling, they provide data on all cells present in the sampled tissues, including less well studied cells such as vascular endothelial cells [3]. Furthermore, transcriptomes obtained through RNA-Seq provide a broader dynamic range than those from microarrays, allowing more precise quantification of the degree to which a gene is expressed in a cell[9]. The major methodological advance of EWCE lies in taking advantage of this improved dynamic range and data availability, in that we are able to use *all* of the genes expressed within a cell type to determine enrichment. This yields a significant improvement in capabilities relative to methods focusing on a small set of selected cell type markers.

Here, we have used EWCE with single cell mouse brain transcriptome data to obtain significant interpretative advances with two important data sources: *(1)* human disease associated genes (2) transcriptomic datasets from post-mortem human brains of diseased and control patients. These applications inform on the primary and secondary cell types involved with brain disorders. The EWCE method is a robust approach, detecting consistent cellular signatures across transcriptome datasets from seventeen Alzheimer’s brain regions and two independent Autism datasets. The EWCE method enables diverse sets of omic data to be integrated and can be applied to a wide range of biological problems in metazoan organisms.

## Methods

### Summary of Expression Weighted Cell-type Enrichment (EWCE) method

The EWCE method takes two arguments: (1) a target gene list of length *n* denoted as *T,* and (2) a background set of genes, indexed by *B*. The objective of the method is to determine the probability that the genes in T have higher expression in a cell type than can be assigned to random chance. To find this we need the probability distribution of average expression in the cell of interest amongst gene lists of length *n.* Assume that one wanted to test the enrichment of astrocyte genes in T. From single-cell transcriptome data we know the expression level of every gene in astrocytes. First we calculate the average expression level in astrocytes of each gene in T. We then randomly sample 100000 gene lists from the background gene set, each with length *n.* The probability distribution is then is estimated from the average level of expression in astrocytes in each of these random gene lists. We have released the *EWCE* package through Bioconductor which can be used to run the method on novel datasets. The method is explained in greater detail below.

### Processing of single cell transcriptome data

Raw cell type mRNA expression data was downloaded from the Linnarsson lab webpage (data annotated as being from 17^th^ August 2014)[3]. Throughout the methods section we refer to this as the Single-Cell Transcriptome (SCT) dataset. The data downloaded provides annotations of the cell type (e.g. ‘astrocyte’, ‘interneuron’, ‘oligodendrocyte’) and sub-cell type annotations (e.g. Int1 [‘interneuron type 1’], Int2 [‘interneuron type 2’], Int15 [‘interneuron type 15’], Peric [‘Pericyte’], Vsmc [‘Vascular Smooth Muscle Cell’]) which are expected to be analogous to either particular cell types (e.g. basket cells) or maturational stages (e.g. differentiating oligodendrocytes). We refer to these as ‘cell types’ and ‘sub-cell type’ annotations. The annotations were not known prior to sequencing and were instead determined using the backspin algorithm as described in the manuscript associated with the dataset. The cells were sampled from two brain regions, SS (somatosensory cortex) and hippocampus CA1: the pyramidal neurons are divided at a cell-type level into these groups. The cell types analyzed in this study are specified and described in Supplementary Table 1.

The dataset contains data from *w* cells associated with *k* sub-cell types. Each of the *k* sub-cell types can be associated with a numerical index from the set {1,..,*k*}. The cell type annotations for cell *i* are stored using a numerical index in *m_i_*, while sub-cell type annotations are stored in *l_i_*. For the EWCE algorithm, we need a single number to describe the relationship between a gene and a sub-cell type. When the number of cells from the cell-type indexed by *c* is given by *N_c_* we calculate the mean level of expression for gene *g* as

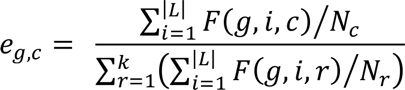

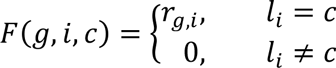

where *r_g,i_* is the expression of gene *‘g’* in cell ‘ *i*’ as described in the data file downloaded from the Linarsson lab website. Because *e_g,c_* is independent of the overall expression level of a gene, it is desirable to drop genes with very low expression levels, as a small number of reads in one cell can make the gene appear to be a highly specific cell marker. We thus drop all genes for which 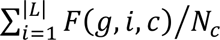is not greater than 0.2 for at least one sub-cell type.

For some of the later analyses (when testing enrichment in genetic susceptibility genes) the values for expression in cell types rather than sub-cell types are used (definition stated above). For this, the values for the expression matrix *E* are first calculated separated for sub-cell types (I.e. S1 Pyramidal cells from Layer 6b) then summed to get the values for grouped cell types (i.e. S1 Pyramidal cells). To get the expression level of gene *g* for the cell type indexed by *e*:

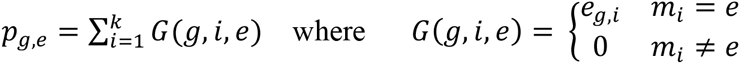

For the remainder of the methods section all the formula’s are provided using *e_g,c_* (the measure of expression in subcell types) but *p_g,e_* should be used instead if it is desired to apply the method to full cell types instead.

### Bootstrapping enrichment within gene lists

The background set used for EWCE analysis depends on the analysis being performed. For gene set (but not transcriptome) enrichment the background gene set is comprised of all genes which have orthologs between human and mice—including those in the target list—but excluding any which were not detected in the SCT dataset. For transcriptome enrichment analysis the background set has an additional restriction relative to simple gene set analysis, in that the background genes must also be expressed in the disease transcriptome dataset. Human genes are converted to mouse orthologs based using Biomart.

The proportion of expression in each cell type is calculated as a matrix for each gene, then summed to get total expression in each cell type across the whole gene list. Thus for a gene list indexed by *X* we calculate the average expression in the sub-cell type indexed by *c* as:

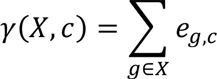

This calculation is then repeated for 100,000 randomly generated gene lists, having the same length as the target gene list, with the genes randomly selected from the background gene set. Sampling of random gene sets is done without replacement. When the target gene list has length *n*, *D_j_* denotes the *j*^th^ set of *n* indices for bootstrapping genes, with *j*∈ {1, … ,100000}. The probability of cellular enrichment is then calculated based on the number of bootstrapped gene lists with higher cell type specific expression than the target list

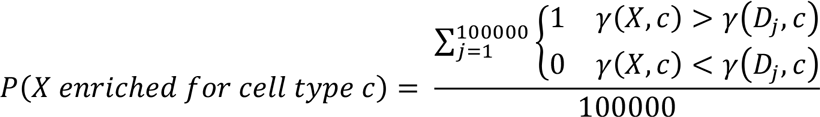

Where probabilities are stated for gene list enrichments, all p-values stated are adjusted for multiple testing. Multiple testing corrections are done separately for the results shown in Figure 2, Figure 3a, 3b and 3c. In the section on schizophrenia transcriptomics, for ease of reading we report p-values for cell-types though the analysis was done on the level of sub-cell types: the p-values reported are for the subcell-type with the most significant relevant enrichment. The fold enrichment *φ(X, c)* is calculated as the expression in the target gene list divided by the mean level of expression in the bootstrap samples

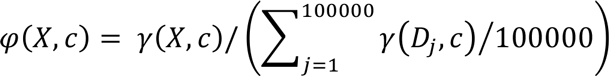

Throughout the paper the number of standard deviations which *γ(X, c)* falls from the mean of *γ(D,c)* is used as a measure of significance. We denote this value for gene list *X* as 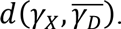

**Figure 2.**
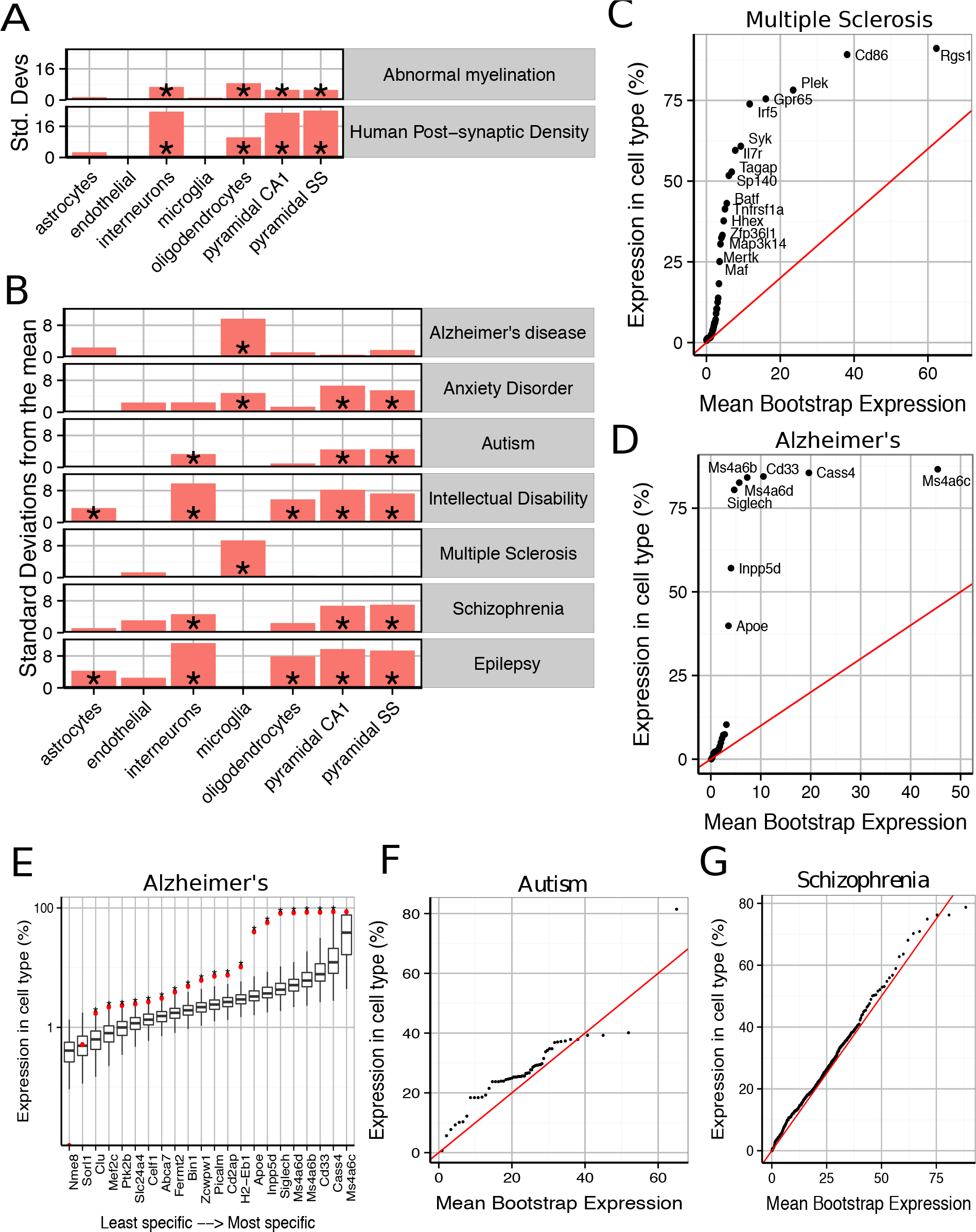
Susceptibility genes for major human brain disorders show distinct cell type enrichments. *(A) Two gene sets with a strong prior expectation for cell type enrichment—the human postsynaptic density, and genes with Human Phenotype Ontology annotations for abnormal myelination—were detected using bootstrapping to have higher expression in neurons and oligodendrocytes respectively.* *(B) Bootstrapping tests performed using the EWCE method show that seven different classes of brain disorder show enrichment in particular cell types.* *(C) Multiple Sclerosis associated genes are strongly enriched for microglial expression. This plot shows that this is not just a property of a few genes, but instead almost every single gene shows higher levels of expression in microglia than would be expected by chance. The plot shows the actual level of expression of the susceptibility gene, against the mean expression level of the i^th^ most expressed gene in a bootstrapping analysis of lists of 19 genes. If microglial expression in MS genes was randomly distributed, the genes would be expected to fall along the red line.* *(D) All Alzheimer’s disease susceptibility genes are more enriched for microglial expression than expected by chance.* *(E) Bootstrap distributions of expected microglial expression levels of Alzheimer’s disease genes. Red dots mark the expression level of the susceptibility genes, while the associated boxplots denote the expected expression level of the i^th^ most expressed gene, in a list of 19 genes, as determined using bootstrapping. Asterisks behind the red dots denote that the gene has higher expression in the cell type than expected by chance (p<0.05).* *(F) Though hundreds of genes are associated with Autism, they are all found to show a moderate increase in expression in pyramidal neurons.* *(G) The hundreds of genes associated with Schizophrenia are found to show a moderate, but highly significantly, increased expression in pyramidal neurons.*

**Figure 3.**
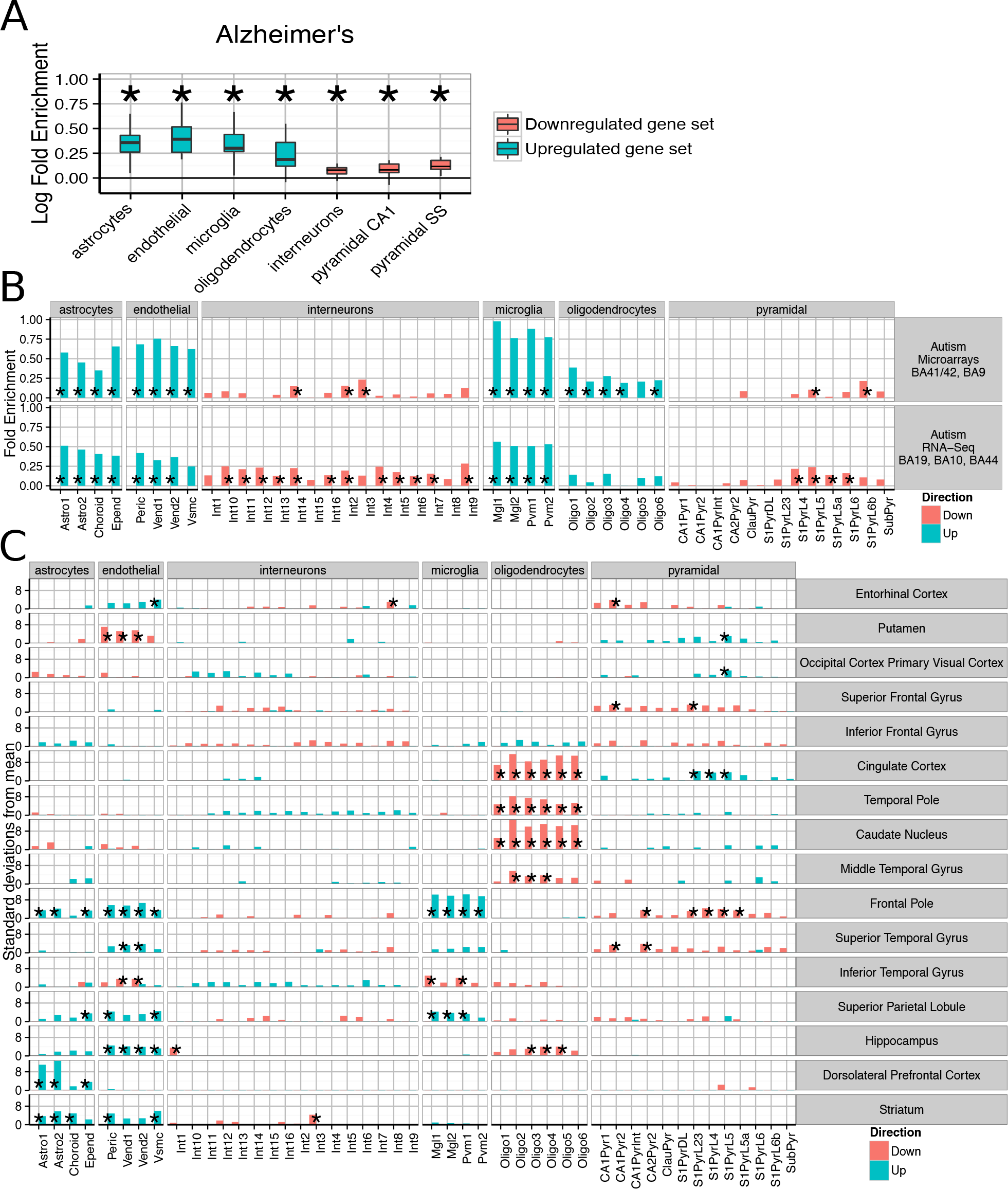
Post-mortem transcriptomes from patients with Alzheimer’s disease, Autism and Schizophrenia show distinctive cellular phenotypes. *(A) Consistent fold enrichments were found for each cell type across fourteen cortical and three subcortical brain regions of Alzheimer’s patients. The boxplots mark the distribution of fold enrichments for that cell types seen across all the brain regions examined. Asterixes mark that the fold enrichment for each cell type that was found to be significantly non-zero with p<0.05.* *(B) Two independent autism studies show the same cellular phenotypes, including upregulation of glial cells and downregulation of neurons. Asterisks mark those cell types found to be significantly differential with p<0.05 after BH correction over all groups.* *(C) Cellular phenotypes in Schizophrenia are regionally dependent but cluster into groups, with a number of regions including the cingulate cortex and temporal pole showing downregulation of oligodendrocyte genes while the prefrontal cortex exhibits upregulation of endothelial and astrocyte genes as well as downregulation of deep pyramidal neurons in the anterior region. The analyses shown are based on an integrative analysis of six independent studies, though not all brain regions featured in all studies.*

### Bootstrapping with controls for transcript length and GC content

Enrichments found in gene sets from genetic studies have been shown to be biased by gene characteristics including transcript length and GC content [10]. To account for this, for the analyses associated with Figure 2 we used a semi-random form of gene selection which controlled for these two properties. For each human gene list, we first obtained the ensembl ID’s associated with the gene symbols through Biomart and then ran a second query to obtain the transcript lengths and GC content associated with each ensemble ID. Where multiple transcript lengths were associated with a single HGNC gene we took the mean value. The deciles of gene size and GC content were calculated over the set of genes expressed in the SCT dataset (after dropping those with low expression levels as described above). The two sets of decile values were used to define a grid, and each gene assigned to a position within the grid based on it’s transcript lengths and GC content. To run a bootstrap analysis on a particular target list, 100000 random lists were constructed with equal length to the target list. Gene *i* in each random list was selected from the same grid square as gene *i* in the target list.

### Disease gene association lists

The disease gene associations were curated from the literature, being largely based on the most recent and authoritative studies. The sources are shown in and the genes comprising each list are in Supplementary Table 2. References associated with the disease lists are listed here: Abnormal myelination[11], Human post-synaptic proteome[12], Alzheimer’s disease[13; 14], Anxiety disorders[15], Autism[16], Intellectual Disability[16], Multiple Sclerosis[17], Schizophrenia[18; 19; 20; 21; 22]. Genes associated with Intellectual Disabilities and Epilepsy were also sourced by finding all genes associated with ‘Intellectual Disability’ and ‘Seizure’ Human Phenotype Ontology terms using the HPO Browser (http://compbio.charite.de/hpoweb). Enrichment probabilities were corrected using the Bonferroni methods.

### Application of EWCE to human disease transcriptome datasets

The transcriptome datasets used in the study were all obtained from publically available sources [23; 24; 25; 26; 27; 28; 29; 30; 31] that are detailed in Supplementary Methods. The methods used to calculate differential expression for each study are also detailed in supplementary methods. The R package *Limma*[32] was used for all tests of differential expression. For all Autism and Schizophrenia studies differential expression was determined for a diagnosis of disease. For Alzheimer’s disease, differential expression was calculated for Braak score. For each study, genes were ordered based on the t-statistic. The 250 genes with the largest positive t-statistics were taken as the upregulated gene set, and the 250 genes with the largest negative t-statistic were taken to be downregulated. Where multiple probes targeted the same gene these were dropped after selected the 250 genes, and the length of the random lists set to have the length of the number of unique genes.

For the Schizophrenia analysis, data from multiple independent studies was available for a number of brain regions. Superior temporal gyrus came from two studies[27; 29]. Middle temporal gyrus was based on two studies[27; 28]. Hippocampus was based on three studies, including one Bipolar cohort[26; 27]. Frontal pole was based on two studies[27; 31]. Dorsolateral prefrontal cortex was based on five studies, including one Bipolar cohort[26; 27; 28; 30]. Cingulate cortex was based on three studies, including samples from anterior and posterior areas[27; 28].

To merge these schizophrenia datasets together the EWCE method was extended as follows. Standard cell type bootstrapping was done for each individual study and the cell type expression proportions for each bootstrap sample was stored as a matrix, with a row for each of the 100000 bootstrap replicates and a column for each cell type. For each individual study being merged, the bootstrap output matrices were summed to form a consensus estimated distribution of random cell type expression proportions. The cell type proportions for the target gene list in each individual study were summed. Calculation of p-values and fold enrichment was then performed as for an individual study, as described above.

## Results

We first sought to confirm that the method detects expected cell type enrichments (Figure 2a). We first gene list we used were the 1461 genes associated with the human cortical postsynaptic density (hPSD) [12]. As expected the hPSD was significantly associated with cortical pyramidal neurons (Bonferroni corrected p<0.00001, 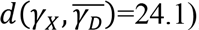, interneurons (p<0.00001, 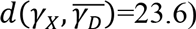 and hippocampal pyramidal neurons (p<0.00001, 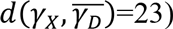. Enrichment was also found for oligodendrocytes (p<0.00001, 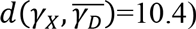, which is likely explained by myelin contamination of the protein samples (amongst the most oligodendrocyte enriched genes in the hPSD list are many of the major protein constituents of myelin including Mbp, Mag, Mog, Pllp and Cnp). The enrichment can also be partially attributed to precursor cells of oligodendrocyte have also been shown to develop post-synaptic densities which are expected to be comprised of many of the same proteins as in neurons[33].

We then examined 185 genes which have Human Phenotype Ontology annotations for abnormal myelination with the prior hypothesis that they would be enriched for oligodendrocyte genes. The majority of the genes are associated with rare neurological disorders in which patients/families have been shown to exhibit either demyelination or absence of myelinated fibers. EWCE analysis confirmed that oligodendrocytes are the cell type most enriched amongst these genes (p<0.00001, 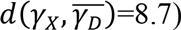. All classes of neurons were also found to be enriched with p<0.00001 suggesting that in many cases the myelination deficit is secondary to a neuronal change.

### Cell enrichments in human disease susceptibility genes

We then tested for cell type enrichments in susceptibility genes for seven major brain disorders: Alzheimer’s disease, Anxiety disorders, Autism, Intellectual Disability, Multiple Sclerosis, Schizophrenia and epilepsy (gene lists used shown in Supplementary Table 2). Two of the disorders were found to show strong evidence for being primary microglial disorders (Alzheimer’s, p<0.00001, 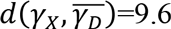 and Multiple Sclerosis, p<0.00001, 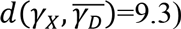. We denote the number of standard deviations from the bootstrapped mean as 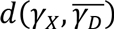. It has previously been noted that a number of Alzheimer’s disease susceptibility genes have high levels of expression in microglia[34], and so we tested whether the enrichment seen here is a result only of expression in those few genes, or whether all susceptibility genes have higher levels of expression than expected by chance. We found that evidence strongly supports the latter hypothesis for both Alzheimer’s and Multiple Sclerosis (Figure 2c—e): for Alzheimer’s, every gene except Cass4 was found to have higher expression in microglia than expected by chance (Figure 2e). No other cell type was found to be significantly enriched for either of these disorders.

Schizophrenia and Autism were found to be the only exclusively neuronal disorders, and both had their strongest enrichment in pyramidal rather than interneurons. Schizophrenia associated genes were enriched for all three classes of neurons with p<0.00001 but for cortical pyramidal neurons 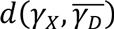=7 while for interneurons 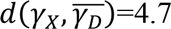. For autism the enrichment for interneurons was close to the significance threshold (p=0.024 and 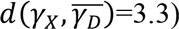 indicating that it may be secondary to the enrichment of pyramidal neuron genes (p=0.00126 and 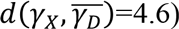. The enrichments for both of these disorders were again found to affect all genes throughout the lists, rather than a subset of strongly cell-type specific genes (Figures 2f--g). Anxiety disorders were also found to be enriched for pyramidal neuron genes (p<0.00001, 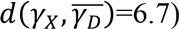 but also showed evidence for a microglial involvement (p<0.00001, 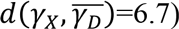.

Interneurons were found to be the most enriched cell type for intellectual disabilities (ID) (p<0.00001, 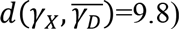 and epilepsy (p<0.00001, 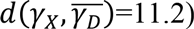. Both disorders also showed enrichment for pyramidal neurons but to a lesser degree (for cortical pyramidal neurons 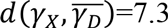for ID and 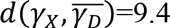 for epilepsy). Astrocytes (p=0.003, epilepsy; p=0.034, ID) and Oligodendrocytes (both p<0.00001) were also found to contribute to both of these conditions. We note one gene in particular, Aass, which is markedly specific to astrocytes and is associated with both seizures and intellectual disabilities.

### Robust cell enrichments in post-mortem disease transcriptomes

We next sought to ascertain whether the method could be used to describe the cellular nature of disease phenotypes found in post-mortem brain samples. Many transcriptome studies have been performed for major brain disorders, in an effort to cast light on the pathological basis of the conditions. We reasoned that because our method utilizes genome-wide data to define ‘set membership’ in a quantitative and brain specific manner, it may be more robust and relevant than GO enrichments at detecting the hidden variables underlying brain diseases.

To perform this analysis, we first perform a standard differential expression analysis and rank order the affected probes by t-statistic, then take the 250 most upregulated, and 250 most downregulated genes. We then perform an Expression Weighted Cell type Enrichment (EWCE) analysis, wherein the random samples are obtained by reordering the ranked list 100,000 times (see Figure 1a for workflow).

The pathological characteristics of Alzheimer’s disease are relatively well understood, with inflammatory gliosis and synapse loss becoming more acute as the disease progresses. We applied the method to an Alzheimer’s dataset that examined changes in fourteen cortical and three non-cortical regions, with between 51—70 samples per region. Differential expression was calculated for genes whose expression is affected by increases in Braak score. Across all the brains tested, a consistent cell enrichment signature was detected (see Figure 3a). We calculated the fold-enrichment for each cell type in each region, for both up- and down-regulated genes, and applied Bayesian estimation to determine whether an actual fold enrichment of zero falls within the 95% Highest Density Interval (HDI): for each of the cell types, we found this to not be the case. Interneuron and pyramidal neuron genes were found to be enriched amongst those which were down-regulated, while oligodendrocytes, microglia, astrocytes and endothelial cells showed evidence of up-regulation. The enrichments detected are directionally consistent with the known pathology of the disease[35; 36], strongly supporting the notion that this is a powerful and robust technique for determining hidden variables in transcriptome data.

Having validated the method’s ability to detect known pathological phenotypes, we sought to apply the method to two disorders that are less well characterized: Autism and Schizophrenia. Two publically available transcriptome datasets were used for Autism[24; 25], with one study using samples from areas BA41/42, BA9, BA19, BA10 and BA44. The two studies were analysed separately, and within each study differential expression was tested over all cortical regions. Remarkably, we again found a consistent cellular enrichment signature across the two studies (Figure 3b). We emphasise that these were two independent studies, performed by different laboratories, on different cortical regions, with one study using RNA-Sequencing and the other Illumina microarrays.

Both Autism studies, like the Alzheimer’s disease studies, indicated that the disease processes affect every major cell type in the brain. As in Alzheimer’s disease, interneuron and pyramidal enrichments were found in down-regulated genes, while enrichments of glial and endothelial cells were present in the up-regulated gene sets. The cellular changes expected to correspond to decreased expression of neuronal transcripts is unclear, with no consensus in the literature about how neurons are affected in autistic patients[37; 38; 39]. Up-regulation of astro- and microglial genes, which indicates activation in those cell types, is broadly supported by past studies which have shown elevated expression of glial marker genes, increased cell densities and altered morphologies for both cell types[40; 41; 42; 43]. The finding that endothelial genes are up-regulated could be related to the decrease in cerebral blood flow seen in temporal and frontal cortices of autistic patients[44].

We next extended the study to Schizophrenia. We utilised data from six independent transcriptomic studies, providing data for many brain areas. Four of the studies included samples from the dorsolateral prefrontal cortex, while other brain regions including hippocampus and cingulate cortex were covered by at least two of the datasets. To maximize the utility of these replicate studies, the cell type bootstrap data was summed across each independent cohort allowing pooled estimates for cellular changes. As was expected based on the disease literature, regional changes were found to be divergent. Some regions (e.g. the primary visual cortex) were found to show no significant changes, while the most pronounced enrichments were seen in the prefrontal and cingulate cortices.

Those regions showing alterations were found to cluster into two groups: (1) those with decreased oligodendrocyte expression and upregulation of pyramidal neuron genes; (2) increased astrocyte and/or endothelial expression. All samples from the prefrontal cortex fell into the second cluster. Four regions fall outside of these clusters and show few/no significant changes: the primary visual cortex, putamen, superior and inferior frontal gyrus. While we showed in Figure 2b that schizophrenia is genetically associated only with neurons, the primary effects appear to be in astrocytes, endothelial cells and oligodendrocytes. A number of regions do show changes in neuronal transcripts, including downregulation of a Somatostatin and Neuropeptide-Y expressing interneuron in the hippocampus (p<0.0266, 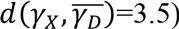, but these changes are less consistent across regions.

One of the most significant changes was in the dorsolateral prefrontal cortex (BA46), with an enrichment of astrocyte genes 12.7 standard deviations from the bootstrapped mean (p<0.00001, 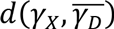=12.7. Significant up-regulation of astrocyte genes (after Benjamini Hochberg correction) was also seen in Striatum (p=0.00001, 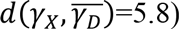, Frontal Pole (p=0.004, 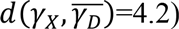 and the Superior Parietal Lobe (p=0.037, 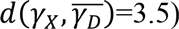. Six regions showed significant up-regulation of endothelial cells in Schizophrenia, including three of those regions which also had up-regulated astrocytes. The most significant effects were seen in the Frontal Pole (p<0.00001, 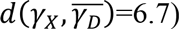 and Striatum (p<0.00001, 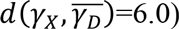. Also affected was the Superior Parietal Lobule (p=0.00992, 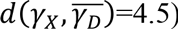, the Hippocampus (p=0.0031, 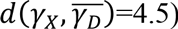, the Entorhinal cortex (p=0.018, 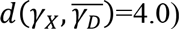 and Superior Temporal Gyrus (p=0.013, 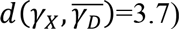. The Frontal Pole, also known as the Anterior Prefrontal Cortex, fits within this cluster but also showed a very strong upregulation of microglial genes (p<0.00001, 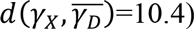 and downregulation of pyramidal neurons (p<0.0009, 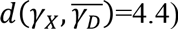.

The second set of brain regions affected in schizophrenia show totally distinct phenotypes from those described above. Astrocyte and endothelial expression appears normal, while highly significant down-regulation of Oligodendrocyte genes was found in the Caudate Nucleus (p<0.00001, 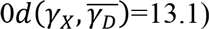, Cingulate Cortex (p<0.00001, 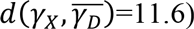, Hippocampus (p<0.0057, 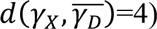, Middle Temporal Gyrus (p<0.0001, 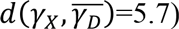. and the Temporal Pole (p<0.0001, 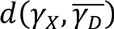=8.2). For four of the five regions the most significant change is in the pre-myelinating (type 2) oligodendrocytes, which are understood to be mid-way through the stages of oligodendrocyte maturation.

## Discussion

Using the EWCE method, we have shown that single cell transcriptome data can be integrated with genetic susceptibility data or tissue transcriptome data to identify cell types involved with disease. Using lists of genetic susceptibility data with single cell transcriptome data that define cell types, we could identify specific cell types that are the likely primary targets of the genetic susceptibility. In a separate analysis, using lists of genes from postmortem transcriptomes, we found that a broader range of cell types were affected, indicating that the cellular pathology of the disease extends from the cells affected by the primary genetic susceptibility to a wider set of “secondary” or “reactive” cell types.

For seven different brain disorders, the EWCE method was used to identify the putative primary cell types affected by genetic susceptibility. Consistent with current models, pyramidal neurons were the cell type most associated with schizophrenia and autism genes. A primary role of microglia in multiple sclerosis is also consistent with primary pathology in the immune system[45]. The association of Alzheimer’s with microglia also builds on a body of existing work on the role of that cell type in the disease[34], but goes further in suggesting it is the primary focus of genetic risk. The identification of anxiety disorders as having a microglial/immune component was surprising and may be relevant to the reported prevalence of anxiety disorders amongst those suffering from immunological disorders[46]. For this later association we caution that unlike the gene sets used for all other disorders, the gene set for Anxiety disorder was generated through a convergent functional genomics approach, which was not solely informed by human genetic studies.

Intellectual disabilities and epilepsy were both found to be most strongly associated with interneurons. Past studies strongly support the finding that interneurons are the causative cell type underlying seizures [47; 48]. Other cell types, including astrocytes and oligodendrocytes were also found to contribute to the etiology of these disorders; this is supported by evidence showing that a human disorder featuring intellectual disabilities and seizures can be attributed to cell-specific mutations in astrocytes [49]. The shared cellular enrichment profiles between ID and epilepsy explains why there exists far high rates of comorbidity between individuals with the two disorders: around 26% of individuals with ID suffer from seizures compared to only 0.4—1% in the general population [50]. Noting the stronger enrichment for interneurons over pyramidal neurons in epilepsy and ID, the findings also suggest an explanation for why individuals with autism are significantly more likely to have seizures if they also have ID[51]: while a mutation affecting synapses in pyramidal neurons may cause Autism, our data suggests that if the mutation affects a gene that is also highly expressed in interneurons this would increase the chance of seizures and ID.

Once denser sequencing of interneurons has been performed from a wider set of brain regions it may be possible to identify specific interneuron populations associated with distinct types of seizures, as well as particular cognitive deficits. The ability to precisely distinguish affected subtypes of interneurons may however require a modification to the method: at present four of the five disorders which are stated as affecting one population of neurons, have significant enrichments for all three neuron types. As neurons share many genes, and vary only in limited subsets and graduated expression levels, it may be that one can only distinguish between them by considering the single most enriched category. The limitations of the method can be seen by considering the case of autism, which shows it’s strongest enrichment in pyramidal cells but is also enriched in interneurons to a lesser degree. These results could support either of two hypotheses: that autism is a primary pyramidal neuron disorder, or that it is a disorder of broad neuronal dysfunction. An extension to the EWCE method that penalizes neuronal subtypes for absence of expression when a gene is expressed in similar cells could potentially resolve this issue.

Contrasting the results of EWCE from genetic susceptibility data with transcriptomes from diseased tissue suggested that the cellular pathology spreads from the primary affected cells to secondary cells. These putative secondary changes appear to extend between classes of cells. For example, the genetic susceptibility of Autism and Schizophrenia appears to primarily impact neurons, yet both show evidence for secondary endothelial disruption. Interestingly, both disorders have been shown to have decreased cerebral blood flow [44; 52; 53] although it is unclear whether this is related to up-regulation of endothelial genes. Though mechanisms are well established for how synaptic activity alters blood flow across brief time scales[54] we are unaware of any studies investigating how persistent mutational changes in the level of synaptic/neuronal activity alters the vascular system. Based on the results we have found here, we suggest that understanding which pyramidal neuronal properties need to be altered (through mutation) to induce secondary transcriptional enrichments of endothelial and/or glial genes could cast new light on the polygenic nature of Schizophrenia and Autism.

Implementation of EWCE in mouse models of human disease could underlie a new approach to studying brain disorders. Once the transcriptomic cell type enrichments are determined for a disease, conditional mouse models (carrying cell type specific mutations) could then be validated or rejected based on whether they recapitulate some or all of the disease associated secondary effects. One problem with this approach is that a range of distinct changes could result in the same transcriptomic alteration (for instance, down regulation of interneuron genes could be caused by either decreased cell density, altered cellular state, or decreased synaptic connectivity). The directionality and regional specificity of transcriptional phenotypes should however be able to act as a guide to narrow down the nature of the cellular changes. Even without more extensive follow-up EWCE could heighten confidence in the biological validity of disease models for which only behavioral similarities to human disease could otherwise be shown.

Three limitations with the current study are that the cells were obtained from mice, they were immature and only from CA1 and somatosensory cortex. Using human rather than mouse data may make a significant difference for disease enrichments: one study which compared expression profiles of cell types between humans and mice found that as few as 52% of genes identified as being astrocyte-enriched in mice, were also found to be astrocyte-enriched in humans[55]. Developmental age is likely to be important since the Braincloud dataset shows that significant transcriptional changes occur in human brain tissue between the pre- to postnatal period[56]. The few anatomical regions so far studied also limits the range of diseases which can currently be analysed. For instance, while a suitable number of genome wide significant genes are known for Parkinson’s disorder[57], the known association of that disorder with the Striatum lead us to avoid testing for enrichment in those genes.

In the coming years the quantity of single cell data that is available will increase and the utility of the EWCE method is expected to expand commensurate with this. The single cell transcriptome dataset used for this study comprised 3005 cells from the cortex and CA1 of mice aged p21--31. With greater depth of cellular sequencing, it may become possible to detect changes in more specific populations of cells. This is potentially of greatest importance for interneurons for which the data presently available is sparse—of the 1314 cells sequenced from CA1, only 126 were from interneurons. With over thirty types of interneuron estimated to exist just within CA1 based on morphology, electrophysiology and expression of classical molecular markers[58], the sparse sampling means there is likely to be substantial noise within the current cell type estimates. Presently it is unclear whether diseases are likely to specifically affect such particular neuron types, and an alternative approach could involve testing along branches of the cell lineage tree—i.e. interneurons derived from medial or caudal ganglionic eminences.

We also note that there is no reason why EWCE should be restricted to the study of brain disorders and as sufficient cellular data becomes available the same methodology could be applied to any other disease. Indeed the EWCE method can be applied to other omic gene lists for the purposes of interrogating the relevant cell types. For example, gene lists from mouse phenotyping studies, such as the International Mouse Phenotyping Consortium[59] could be used to identify the cell types underlying specific phenotypes.

## Acknowledgements

This work was supported by the Wellcome Trust and the European Union Seventh Framework Programme under grant agreements n° HEALTH-F2-2009-241995 (“Gencodys” project) and HEALTH-F2-2009-242167 (“Synsys” project).

